# Differential encoding of predator fear in the ventromedial hypothalamus and periaqueductal grey

**DOI:** 10.1101/283820

**Authors:** Maria Esteban Masferrer, Bianca A. Silva, Kensaku Nomoto, Susana Q. Lima, Cornelius T. Gross

## Abstract

The ventromedial hypothalamus is a central node of the mammalian predator defense network. Stimulation of this structure in rodents and primates elicits abrupt defensive responses, including flight, freezing, sympathetic activation, and panic, while inhibition reduces defensive responses to predators. The major efferent target of the ventromedial hypothalamus is the dorsal periaqueductal grey, and stimulation of this structure also elicits flight, freezing, and sympathetic activation. However, reversible inhibition experiments suggest that the ventromedial hypothalamus and periaqueductal grey play distinct roles in the control of defensive behavior, with the former proposed to encode an internal state necessary for the motivation of defensive responses, while the latter serves as a motor pattern initiator. Here we used electrophysiological recordings of single units in behaving mice exposed to a rat to investigate the encoding of predator fear in the dorsomedial division of the ventromedial hypothalamus and the dorsal periaqueductal grey. Distinct correlates of threat intensity and motor responses were found in both structures, suggesting a distributed encoding of sensory and motor features in the medial hypothalamic-brainstem instinctive network.

## Introduction

Electrical stimulation and lesion studies identified the medial hypothalamus as a central organizer of innate goal-directed behaviors associated with defense and reproduction (Malsbury et al., 1977; Kruk et al., 1979; Pfaff and Sakuma, 1979a, 1979b; Canteras et al., 1997; Canteras, 2002; Lin et al., 2011; Yang et al., 2013; Silva et al., 2013). Evidence for its anatomical and molecular conservation across vertebrates (Braak and Braak, 1992; Koutcherov et al., 2002; Kurrasch et al., 2007) suggests that it forms an ancient control center for survival responses that may date back as far as the evolution of bilaterians (Tessmar-Raible et al., 2007; Arendt et al., 2015). Anatomical tract-tracing and cFos mapping studies in rats exposed to cats identified a set of three interconnected medial hypothalamic nuclei, the anterior hypothalamic nucleus (AHN), dorsomedial division of the ventromedial hypothalamus (VMHdm), and dorsal premammillary nucleus (PMD), that together comprise the medial hypothalamic defensive system (Canteras, 2002). The VMHdm occupies a central position in the network because it receives direct input from olfactory and pheremonal sensory processing areas in the cortical and medial amygdala (Swanson and Petrovich, 1998; Choi et al., 2005; Bergan et al., 2014) necessary to detect threat-related stimuli and at the same time provides direct outputs to the dorsal periaqueductal grey (dPAG), considered to be the behavioral and autonomic motor pattern initiator for defensive responses (Blanchard et al., 1981; Bandler and McCulloch, 1984; Fuchs et al., 1985; Graeff, 1994; Swanson, 2000; Canteras, 2002).

Recent work in mice has begun to dissect the medial hypothalamic defensive system at the circuit level. Optogenetic stimulation of neurons in VMHdm (Lin et al., 2011; Kunwar et al., 2015) or their projections to AHN or PAG (Wang et al., 2015) elicited freezing and flight behavior and pharmacogenetic inhibition or genetic ablation of neurotransmission in VMHdm reduced defensive responses to predators (Silva et al., 2013; Kunwar et al., 2015).

Neurons in VMHdm appear to selectively support defensive behaviors elicited by predators, as defensive responses to a conspecific were unaffected by pharmacogenetic inhibition (Silva et al., 2013; but see Kunwar et al., 2015). Similarly, optogenetic stimulation of neurons in dPAG elicited freezing and flight (Deng et al., 2016; Tovote et al., 2016) and pharmacogenetic inhibition reduced defensive responses to predator (Silva et al., 2013).

However, pharmacogenetic inhibition of dPAG did not impair the acquisition of a fear memory formed by exposure to the predator, while VMHdm inhibition did (Silva et al., 2016b), suggesting a role for PAG limited to the expression of defensive behaviors and a broader role for VMH in encoding an internal, motivational state required both for the expression and memory of predator defense (Silva et al., 2016a, 2016b). Data from deep brain electrical stimulation studies in humans appear to support this distinction, with stimulation of VMHdm eliciting feelings of dread, impending doom, and panic (Wilent et al., 2009) and stimulation of dPAG eliciting sensations of being chased (Amano et al., 1982).

A potential role for VMHdm in encoding defensive motivation is also supported by analogy with single unit electrophysiological recordings and calcium imaging in the ventrolateral division of the VMH (VMHvl), a part of the medial hypothalamic reproductive system, in male mice exhibiting aggression toward male intruders (Lin et al., 2011; Falkner et al., 2014, 2016; Remedios et al., 2017). In these studies, VMHvl neurons showed increased firing when exposed to awake males, anesthetized males, or male urine, and the intensity of activation was correlated with the latency and duration of future attacks, demonstrating the encoding of both sensory as well as motivational features of the behavior. The capacity of neural activity in VMHvl to motivate attack behavior was subsequently tested by showing that optogenetic stimulation of VMHvl could drive lever pressing behavior to get access to an intruder (Falkner et al., 2016). Single unit recordings in the VMHdm of behaving animals have not been reported, but by analogy with VMHvl such neural activity should similarly scale with both sensory features of the threat as well as the probability or intensity of the defensive response.

In dPAG, on the other hand, electrophysiological recordings of single units in awake behaving mice exposed to a rat identified two major classes of neurons – flight cells, whose firing increased during flight from the predator, and assessment cells, whose firing increased with decreasing distance from the predator (Deng et al., 2016). These findings demonstrated that dPAG neuron activity encodes not only defensive motor actions (flight), but also sensory aspects of threat distance and intensity (assessment). Together these findings suggest that a simple model in which VMHdm firing encodes threat intensity and dPAG neurons are triggered to produce defensive behaviors when this activity reaches a given threshold is likely to be incorrect. Here we investigated single unit activity in both VMHdm and dPAG in awake behaving mice exposed to a rat to understand whether and how predator defense behavior is differentially encoded in these structures. Our data revealed that both VMHdm and dPAG contain assessment and flight cells, suggesting a distributed encoding of sensory and motor aspects of defense across these structures. However, correlations between defensive behavior and firing rates were different in VMHdm and dPAG, confirming a hierarchical encoding of defense between medial hypothalamus and brainstem.

## Results

Current approaches for understanding the encoding of behavior in the firing of single neurons involve recording single unit electrical activity in awake behaving animals showing robust and repeated responses to an eliciting stimulus over multiple sessions and days. To achieve such conditions we modified our existing mouse predator defense test (Silva et al., 2013) so that the mouse could be transferred each day from its home cage to the testing apparatus consisting of a 60 cm long corridor connected to a larger 25 × 25 cm stimulus chamber via a small opening (**Figure 1A**). At the beginning of each session the electrode connector of the experimental animal was plugged into the recording cable and the animal was returned to its home cage for 20 minutes to stabilize the recording. At the start of the experiment the animal was transferred to the experimental apparatus and allowed to explore during a 10 minute habituation phase. At a moment when the experimental animal was in the corridor compartment a rat was placed into the stimulus chamber and the defensive behavior of the mouse was observed for a further 10 minutes. At the end of the session, the rat was removed and the mouse was allowed to explore for an additional 10 minutes. Quantification of the cumulative dwell time in the apparatus revealed a clear decrease in time spent near the stimulus chamber in the presence of the rat (**Figure 1BC**). Notably, in the presence of the rat the mouse repeatedly carried out a sequence of behaviors in which it moved cautiously toward the stimulus chamber assessment and then turned to initiate a rapid movement (flight) away from the rat (**Figure 1D**). Robust assessment-flight sequences were seen over multiple testing sessions (9.3±3.2 per session; **Figure S1A**; over 13±1.0 sessions per animal, maximum 1 session per day, N=12). No flights were observed during the habituation or post-rat phases (**Figure S1B**).

**Figure 1.**
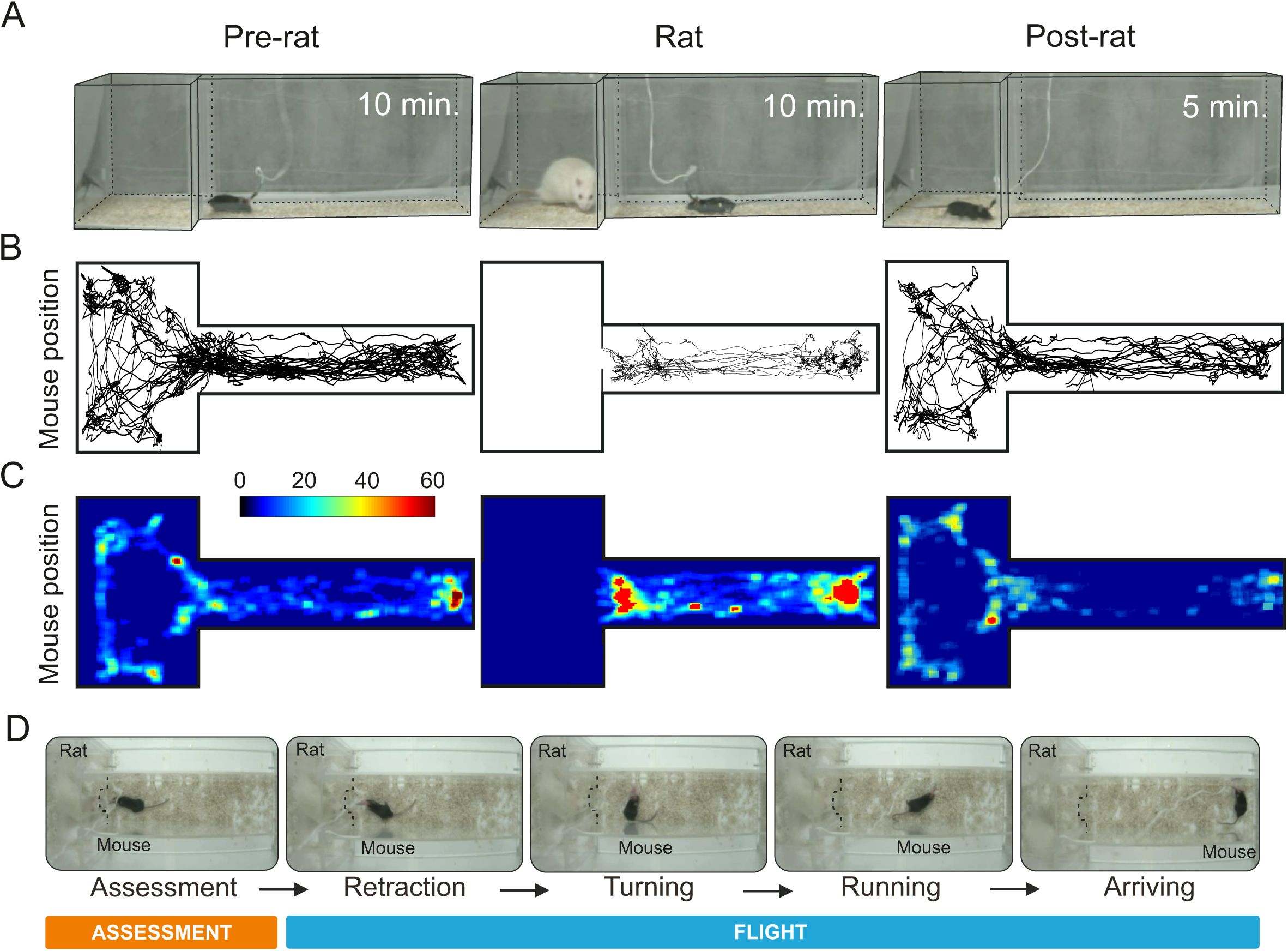
Predator exposure test apparatus. The mouse was transferred to the testing apparatus and allowed to explore the corridor and stimulus chamber first in the absence of the rat, then in the presence of the rat, and then again without the rat. (**A**) Representative example of mouse position during pre-rat (left), rat (middle) and post-rat (right) phases. (**B**) Heat-map showing average cumulative time in the apparatus in the pre-rat (left), rat (middle), and post-rat (right) phases (N = 12). (**C**) Sequence of video frames for a representative mouse executing approach and flight during the rat exposure phase: approach (assessment) and flight (retraction, turning, running, arriving).

Animals were implanted with a 16-microwire electrodes fitted to a movable drive into either the VMHdm or dPAG and allowed to recover for at least two weeks before testing. Electrodes were advanced at the end of each session and putative single units identified before the start of testing using standard spike sorting methods. A total of 305 putative single units recorded in 13 mice across a total of 101 sessions satisfied all criteria and were included in the final analysis. To identify units whose firing pattern was correlated with approach-flight behavior we aligned the firing of units to the start of each flight event across sessions as measured by an overhead videotracking apparatus and assessed correlations statistically.

In dPAG, 43% of single units (30/69) were significantly correlated with approach-flight behavior (**Figure 2B**). The patterns of correlation could be clustered into three types – cells that increased firing during approach (“assessment+” cells, 13%, 9/69; **Figure 2C; Movie S1**), cells that decreased firing during approach (“assessment-“ cells, 12%, 8/69; **Figure 2D; Movie S2**), and cells that increased firing during flight (“flight” cells, 16%, 11/69; **Figure 2E; Movie S3**). One single unit consistently decreased firing during flight and another one correlates with both, approach and flight behavior, but given that it was the solitary unit with this pattern, we did not consider this a robust cell-type. Similar assessment+ and flight cells have been previously reported in dPAG (Deng et al., 2016). As assessment cells increased their firing rate during the approach toward the rat we plotted their average firing rate plotted against distance from the stimulus chamber. This analysis revealed a similar, non-linear relationship between both assessment+ and assessment-cell firing and distance (**Figure 2F**) in which assessment cells moderated their firing only when close to the stimulus chamber. This relationship is consistent with the encoding of threat proximity by dPAG assessment cells. To understand whether flight cells in dPAG might encode motor features of flight we plotted the average firing rate of flight cells superimposed on the animal’s velocity. This analysis revealed that peak flight cell firing occurred before peak velocity was attained (**Figure 2G**) consistent with the encoding of a motor pattern initiator rather than a motor executor function for dPAG flight cells.

**Figure 2.**
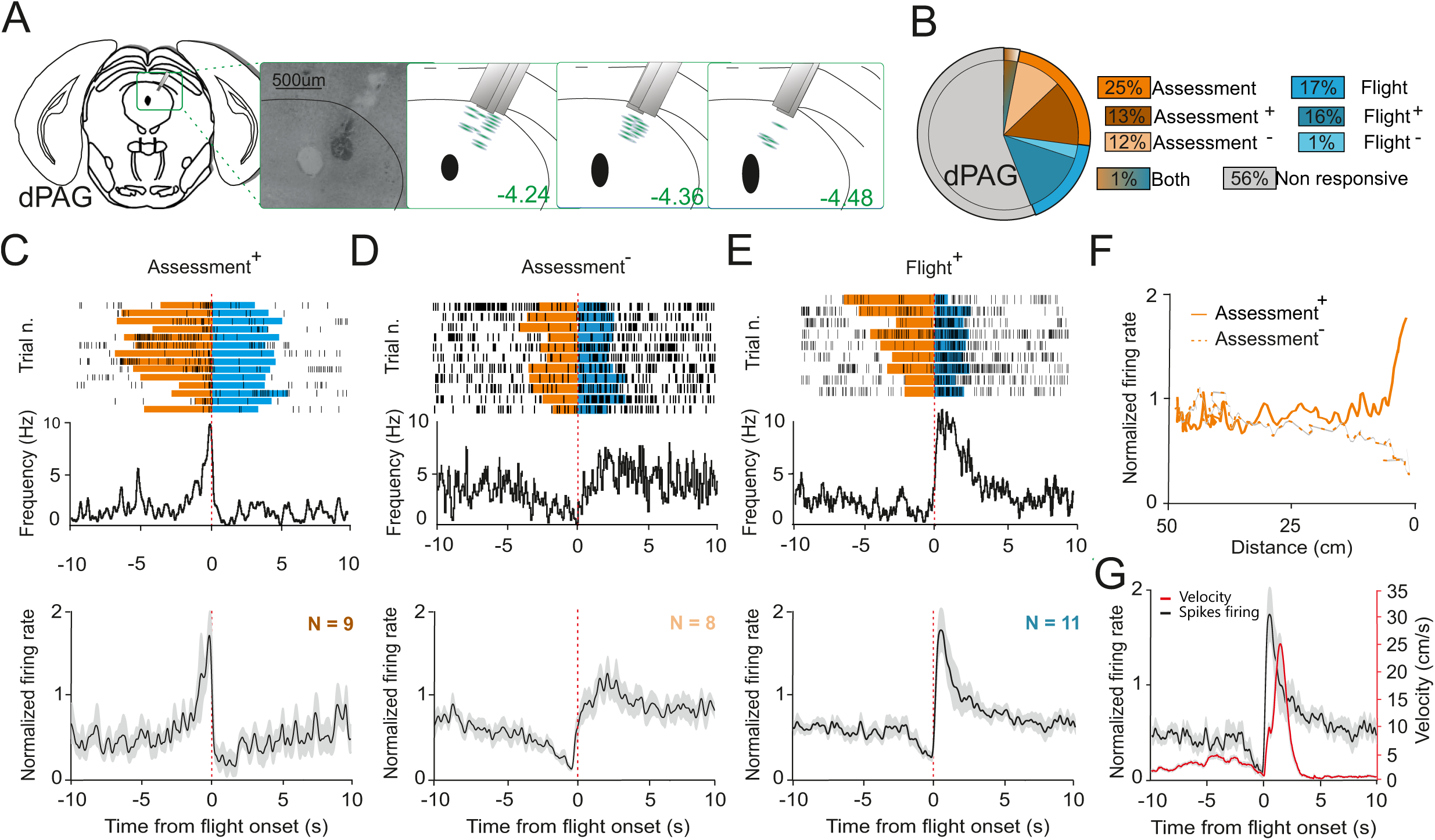
Single unit activity in dPAG during exposure to predator. (**A**) Nissl staining showing representative electrolytic lesion in dPAG and estimated electrode recording sites. (**B**) Population distribution of single units identified in dPAG (25% assessment, 16% flight, 1% both; mice N = 8, cells N = 69). (**C**) Firing frequency of a representative assessment+ cell over trials (top). Normalized average firing rate of all assessment+ cells identified (bottom, N = 9). (**D**) Firing frequency of a representative assessment-cell over trials (top). Normalized average firing rate of all assessment-cells identified (bottom, N = 8). (**E**) Firing frequency of a representative flight+ cell over trials (top). Normalized average firing rate of all flight+ cells identified (bottom, N = 11). (**F**) Normalized firing rates of assessment+ (continuous line) and assessment-(dashed line) cells identified plotted against distance from the rat chamber. (**G**) Average firing activity of flight+ cells and mouse velocity during flight. Time zero represents the flight onset.

In VMHdm, 28% of single units (67/236) were significantly correlated with approach-flight behavior (**Figure 3B**). The patterns of correlation could be clustered into three types – cells that increased firing during approach (“assessment+” cells, 10%, 24/236; **Figure 3C; Movie S4**), cells that increased firing during flight (“flight+” cells, 8%, 19/236; **Figure 3D; Movie S5**), and cells that decreased firing during flight (“flight-” cells, 9%, 21/236; **Figure 3E; Movie S6**). Three single units consistently decreased firing during approach, but given the small number of units with this pattern, we did not analyze this cell-type further. To examine what aspects of approach might be encoded by assessment+ cells we plotted their average firing rate against distance from the stimulus chamber. This analysis revealed a linear relationship between assessment+ cell firing and distance (**Figure 3F**) in which assessment+ cells monotonically increased their firing rate as they approached the rat chamber. This relationship is consistent with the encoding of threat intensity by VMHdm assessment cells. To understand whether flight cells in VMHdm might encode motor features of flight we plotted the average firing rate of flight+ cells superimposed on the animal’s velocity. This analysis revealed that the peak firing of flight+ cells occurred shortly before peak velocity was attained (**Figure 3G**). Using the initiation of flight as a common time point we compared the profiles of firing rates of flight+ cells in dPAG and VMHdm (**Figure 3H**). Peak firing of dPAG flight+ cells occurred on average 220 ms before that of VMHdm flight+ cells. Consistent with an earlier peak firing in dPAG flight+ cells, a comparison of the correlation between flight+ cell firing and velocity revealed a steeper slope for dPAG compared to VMHdm (**Figure 3I**). Although in the absence of data deriving from simultaneous recordings in the two structures we cannot draw conclusions about the relative latency of neuronal recruitment, this comparison suggests that dPAG is more rapidly recruited during flight than VMHdm.

**Figure 3.**
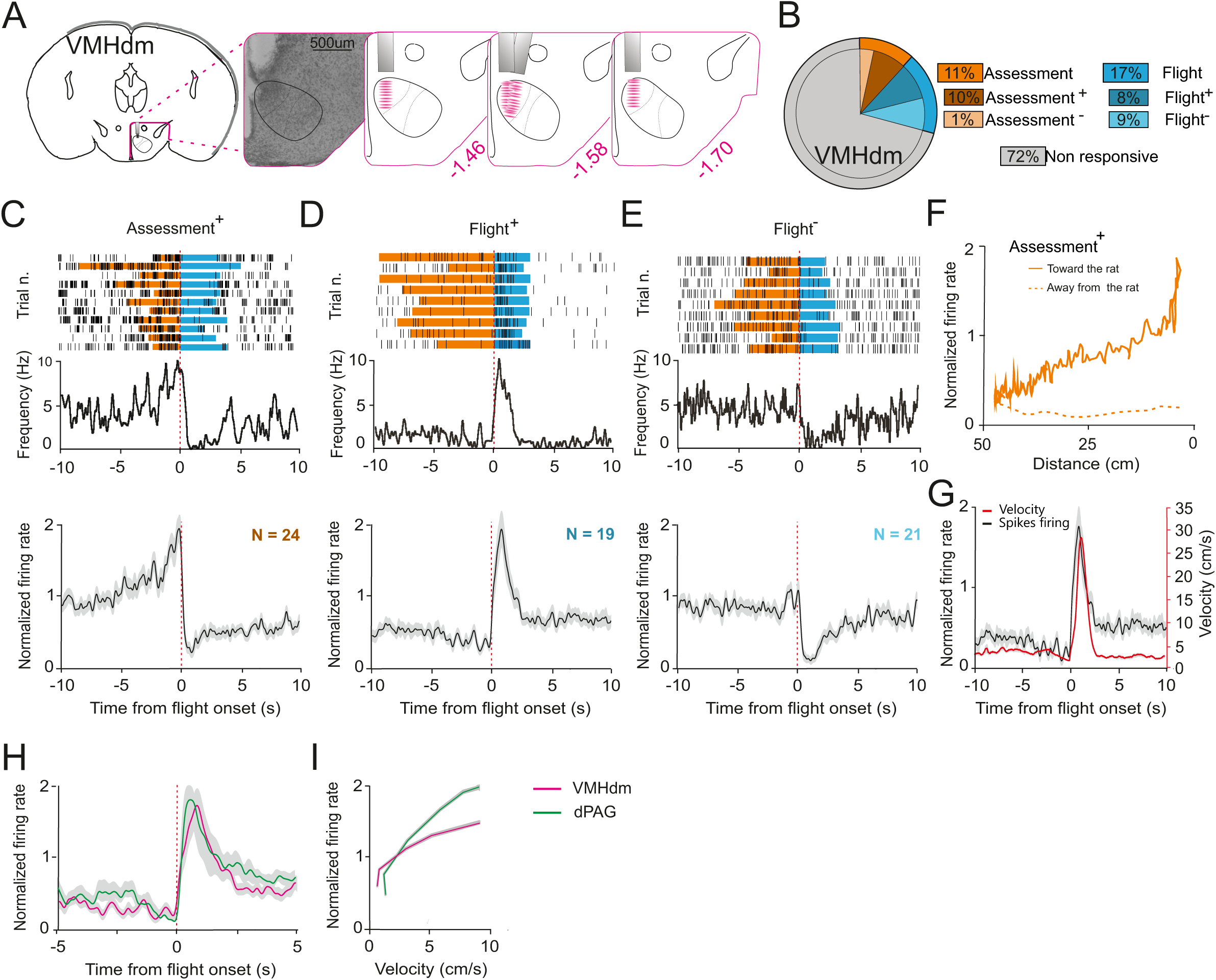
Single unit activity in VMHdm during exposure to predator. (**A**) Nissl staining showing representative electrolytic lesion in VMHdm and estimated electrode recording sites. (**B**) Population distribution of single units identified in VMHdm (11% assessment, 17% flight, mice N = 4, cells N = 236). (**C**) Firing frequency of a representative assessment+ cell over trials (top). Normalized average firing rate of all assessment+ cells identified (bottom, N = 24). (**D**) Firing frequency of a representative flight+ cell over trials (top). Normalized average firing rate of all flight+ cells identified (bottom, N = 19). (**E**) Firing frequency of a representative flight-cell over trials (top). Normalized average firing rate of all identified flight-cells (bottom, N = 21). (**F**) Normalized firing rates of assessment+ cells plotted against distance from the rat chamber. Continuous line indicates firing rate during approach and dashed line indicates firing rate during flight. (**G**) Average firing activity of flight+ cells and mouse velocity during flight. (**H**) Normalized firing rate of flight+ cells in dPAG (green) and VMHdm (magenta) during flight. (**I**) Average firing activity of flight+ cells in dPAG (green) and VMHdm (magenta) plotted against average velocity during flight. Time zero represents flight onset.

Finally, we examined the firing rates of assessment cells in dPAG and VMHdm during the habituation and post-rat phases to determine whether similar increases of firing could be seen when the animal approaches the stimulus chamber in the absence of the rat. Mice showed similar numbers of approaches to the stimulus chamber during the habituation and rat phases (**Figure S1B**), however firing rates of dPAG and VMHdm assessment cells did not increase during approach to the stimulus chamber in the absence of the rat (**Figure S3**). These findings demonstrate that assessment cell firing during approach toward the rat is dependent on predator sensory cues.

## Discussion

We have recorded the firing of 236 putative single neurons in the mouse VMHdm and dPAG during exposure to a live rat. We found two classes of neurons among those that showed firing that was significantly correlated with the defensive behavioral responses of the experimental animals. One class showed firing whose rate was modulated as the animal approached the rat, called “assessment” cells, and the other whose rate was modulated as the animal fled the rat, called “flight” cells. Both classes of cells were found in VMHdm and dPAG, suggesting that both structures encode aspects of threat detection and avoidance consistent with the known requirement of these structures in the expression of predator defense.

However, several differences in the correlations between firing and behavior were evident in VMHdm and dPAG. For assessment+ cells, firing rates increased in an inverse linear manner with distance from the rat in VMHdm, but in a delayed, non-linear manner in dPAG (**Figure 2C, 3C**). This difference suggests that VMHdm is activated earlier during approach to a predatory threat and that it directly encodes threat distance, while dPAG is activated only in close proximity to the predator as threat levels rise beyond a certain threshold. Because VMHdm receives direct projections from the MeApv that encodes predator odor (Canteras et al., 1995; Choi et al., 2005; Bergan et al., 2014), it is reasonable to postulate that the increase in firing of assessment+ cells in VMHdm reflects their direct receipt of predator odor information from upstream olfactory and kairomone processing areas. However, VMHdm is also likely to receive non-olfactory sensory information about the predator from multimodal sensory processing areas of the amygdala, such as the basomedial and basolateral nucleus (Petrovich et al., 1996; Li et al., 2004; Martinez et al., 2011; Gross and Canteras, 2012). Given the highly linear relationship between VMHdm assessment+ cell firing and predator distance, we argue that VMHdm assessment+ cells are directly driven by multimodal sensory information about the threat. This hypothesis was confirmed by our observation that assessment cell firing during the habituation and post-rat phases was not increased as the animal approached the stimulus chamber (**Figure S3**). These data also demonstrate that predatory olfactory cues remaining in the stimulus chamber are not sufficient to recruit VMHdm or dPAG neuronal firing during the post-rat phase, a finding consistent with the failure to recruit cFos activity during predator contextual fear memory (Motta et al.,2009; Silva et al., 2016b), but seemingly at odds with the ability of pharmacogenetic inhibition of VMHdm to inhibit defensive behavior to the predator context (Silva et al., 2016b). However, methodological differences (e.g. housing, handling, and/or training-testing delay) may have reduced the recruitment of VMHdm during predator context recall in our study. Finally, we note that firing activity of VMHvl units during approach to conspecifics is also proportional to distance (Lin et al., 2011; Falkner et al., 2014), although in these studies the small size of the testing apparatus precluded an estimation of the linearity of this relationship.

Given that VMHdm neurons project heavily to dPAG and the hypothalamus provides over 45% of dPAG inputs (Beitz, 1982, 1989; Canteras et al., 1994; Silva et al., 2013; Wang et al., 2015), we speculate that the non-linear relationship with predator distance shown by dPAG assessment cell firing (both assessment+ and assessment-cells) rests in part on their being driven in a non-linear manner by inputs from VMHdm. Although the olfactory and multimodal sensory processing areas of the amygdala do not project directly to PAG (Canteras et al., 1995; Petrovich et al., 1996; Martinez et al., 2011), dPAG assessment cells could also receive multimodal sensory information about the predator from thalamic and collicular inputs (Mantyh, 1982; Beitz, 1989; Vianna and Brandão, 2003; Schenberg et al., 2005; Silva et al., 2013).

For flight cells, firing rates peaked earlier in dPAG than in VMHdm (**Figure 3H**). A delay was also seen in the peak of firing of flight cells relative to the peak in velocity (**Figure 2G, 3G**), with dPAG peaking an average of 2.5 s before peak velocity and VMHdm peaking an average of 0.41 s before. In addition, dPAG flight cell firing increased at a steeper rate relative to the animal’s velocity than VMHdm (**Figure 3I**). Although without data from simultaneously recorded VMHdm and dPAG flight cells we cannot draw definitively conclusions about the relative latency to initiate firing of flight cells in the two structures, the steeper correlation with velocity suggests that dPAG is recruited in a robust manner earlier or more strongly than VMHdm during flight and may be driving it, an interpretation that is consistent with the existence of significant back-projections from dPAG to VMHdm (Mantyh, 1983; Meller and Dennis, 1991; Cameron et al., 1995).

It is not clear why so few assessment-cells were found in VMHdm and why so few flight-cells were found in dPAG. Given that assessment+ and flight+ cells with similar behavioral correlations were found in each structure, it is possible that their relative paucity reflects an absence of local inhibitory collaterals of assessment+ and flight+ cells in VMHdm and dPAG, respectively.

Our findings are consistent with an earlier study in which assessment+ and flight+ cells were described in the mouse dPAG during exposure to a rat (Deng et al., 2016), although that study did not report approach or flight cells with decreasing firing rates, a discrepancy that may derive from the normalization methods used for identifying behaviorally correlated units. Like in the previous study, firing of dPAG assessment+ cells increased only when the mouse approached closely to the rat chamber (**Figure 2F**), suggesting that they might take part in a mechanism to bring dPAG activity to a threshold sufficient to activate local flight cells that signal the expression of escape behaviors when threat intensity is maximal. Interestingly, firing of assessment+ cells is shut off upon flight initiation, suggesting the existence of a negative feedback mechanism from flight to assessment cells, possibly depending on dPAG back-projections. Where does the threat information come from that drives assessment+ cell activity in dPAG? VMHdm assessment+ cells show a more linear response to threat proximity and one possibility is that these cells drive dPAG assessment+ cells to threshold. However, direct optogenetic stimulation of VMHdm-dPAG projections was not able to elicit flight behavior (Wang et al., 2015), suggesting that other dPAG afferents may be required. Inputs from the dorsal pre-mammillary nucleus (PMD), for example, are required for flight responses to predator and these may provide the critical drive for dPAG flight cells to reach threshold (Canteras and Swanson, 1992; Blanchard et al., 2003). At the same time, VMHdm assessment+ cells are likely to be driven by amygdala inputs carrying multi-modal sensory information about the predator.

One of the surprising findings of our study was the presence of flight cells in VMHdm. Their presence demonstrates that the medial hypothalamus encodes both sensory and motor information about defense. It remains to be determined which inputs drive VMHdm flight cells. One possibility is that they are driven by thresholding of local assessment+ cells, a mechanism that would require a similar mechanism to drive flight-cells in VMHdm, a possibility that we consider unlikely, given the paucity of inhibitory interneurons in this subnucleus (Lein et al., 2007). Another possibility is that flight+ and flight-cells in VMHdm are driven by feedback from flight cells in dPAG via prominent feedback projections known to exist between these structures (Mantyh, 1983; Meller and Dennis, 1991; Cameron et al., 1995). The earlier and steeper firing of dPAG compared to VMHdm flight cells supports this hypothesis (**Figure 3I**). Regardless, our data show that medial hypothalamus and its major brainstem target encode both sensory and motor features of defensive behavior, albeit with different correlations to behavior. In particular, we show that information about threat intensity and defensive motor action (flight) coexist already at the level of the medial hypothalamus.

## Materials & Methods

### Animals

All test subjects used were adult C57BL/6 N male mice, 8 weeks old at the moment of the surgery, from the European Molecular Biology Laboratory internal breeding colonies. Predators were adult male SHR/NHsd rats from Harlan Sprague Dawley Inc. All animals were singly housed after the surgery at 22-25 °C under a 12-h light-dark cycle with water and food ad libitum. All animals were handled according to protocols approved by the Italian Ministry of Health.

### Surgery

The stereotaxic surgery (Kopf Instruments) was performed under deep anesthesia (isoflurane 2%). The skull surface was exposed and two stainless steel screws (RWD Life Science) were fixed permanently into the posterior and anterior portions of the skull, to serve as ground and fixation. For extracellular recordings, mice were implanted with a movable bundle of 16 tungsten microwires (23 µm in diameter, Innovative Neurophysiology; Nomoto et al, 2015), into the dmVMH (-0.95 mm posterior to bregma, +0.3 mm lateral, and -5.35 mm ventral to the brain surface) or into the dPAG (-4.1 mm posterior to bregma, +1.18 mm lateral and -2.36 mm ventral to the skull surface, at 26° lateral angle). The bundle was fixed on the skull using dental acrylic (DuraLay Reliance). After surgery mice were injected with carprofen (Rimadyl, 5mg/kg s.c.) for three days to control pain and inflammations. All animals were allowed to recover for at least 2 weeks before any test was performed.

### Behavioral assay

The experimental apparatus (adapted from Silva et al., 2013) was made of transparent Plexiglas and composed of a stimulus chamber (25 × 25 × 25 cm) that was connected by a removable panel with an opening (2.0 wide, 2.0 cm high) to a narrow corridor (12.5 cm wide, 60 cm long, 30 cm high). At the beginning of each session the experimental animal was connected to the acquisition system and monitored for 20 min in the home cage to confirm signal stability before being placed into the experimental apparatus. Each session consisted of 10 min of habituation in which the door with the opening was removed in order to allow the animal to freely explore the corridor and stimulus chamber, followed by replacing the door with the opening and the placement of the rat in the stimulus compartment. Sensory contact between the rat and the subject was permitted for 10 min. Subsequently, the rat and opening panel were removed and the mouse was allowed to explore the corridor and stimulus chamber for another 5 min. Videos were recorded at 30 frames/s to extract position and velocity of the animal (CinePlex Studio, Plexon). Video and neural data were synchronized offline and the behavior was manually scored (CinePlex Editor, Plexon).

### Electrophysiology

During the recording sessions mice were connected to a lightweight head stage (1.03 g., gain 20x, Plexon). The head stage was connected to a 16-channel analog amplifier (gain 50x, Plexon) and neural activity was checked online. If at least one unit was identified, the behavioral paradigm started. Otherwise, the bundle of electrodes was re-adjusted by moving the screw to advance it by 60 µm followed by a waiting period of 24 h before the next recording session. The neural signal was acquired (digitized at 40 kHz, band pass filtered from 300Hz to 8 kHz) and stored using a Neural Data Acquisition System (Omniplex, Plexon) for offline analysis. Spikes were sorted offline (Offline Sorter, Plexon) based on 3D principal component analysis. Units isolation was verified using autocorrelation histograms, and cross-correlation histograms were used to detect units appearing in more than one channel. Three criteria were used to consider the recorded cell as a single unit: 1) signal-to noise ratio >3, 2) stable waveform shape during the recording, and 3) percentage of spikes occurring with inter-spike intervals <2 ms must be <0.1%. At the end of each session the electrodes were advanced 60 µm.

### Physiology analysis

Confirmation of well-isolated single units was performed using NeuroExplorer software. Peri-stimulus time histograms (PSTHs) were computed using custom Matlab (Mathworks) scripts. For each neuron, a mean spike density function was constructed by applying a Gaussian kernel (σ=10 ms). PSTHs were calculated using 10 ms bins. For the mean PSTH, the firing rate for each unit was normalized to the mean frequency of that unit across the whole session and averaged across trials. To define units as “assessment” or “flight” cells they had to show a significant firing rate variation during risk assessment and flight compared with a 2 s randomly chosen baseline time window (Wilcoxon rank-sum test, p<0.001; errors in ±SEM).

### Histology

At the end of the experiment a small electrolytic lesion was made around the tip of the electrode (20 µA, 40 s; Cibertec stimulator). Serial coronal sections (40 µm) of perfused brains (1x PBS and 4% PFA in 1x PB) were cut on a cryomicrotome, stained using Nissl technique (0.1% Cresyl Violet) and visualized under the microscope to verify the electrode placement.

## Acknowledgements

We thank Francesca Zonfrillo, Roberto Voci, Matteo Gaetani, and Monica Serra for mouse husbandry and Angelo Raggioli for laboratory management support. The work was supported by EMBL and the European Research Council (ERC) Advanced Grant COREFEAR to C.T.G.

## Author contributions

All the behavioral and electrophysiological experiments and their analysis were carried out by M.E.M. except for initial exploratory experiments carried out by B.A.S. and K.N. in the laboratory of S.L. M.E.M. and C.T.G. conceived the project, designed the experiments, and wrote the manuscript.

## Supplementary Figure Legends

***Figure S1.* (A)** The number of flights/session in animals implanted with electrodes in dPAG (green) and VMHdm (magenta) were not significantly different. **(B)** Number of approaches ending with and without flight in different phases of the recording session.

***Figure S2.* Summary of single unit firing rates.** (**A, B**) Firing rate of all assessment+, assessment-, flight+ and flight-cells identified in VMHdm and dPAG, respectively. (**C**) Normalized firing rate of assessment+, assessment-, flight+ and flight-cells in dPAG (green) and VMHdm (magenta) during flight onset.

***Figure S3.* Average of single unit firing rates during different phases of predator exposure. (A)** Mean firing rate of all assessment+, assessment-, flight+, and flight-cells identified in VMHdm (**Top row** – pre-rat phase, **middle row** – post-rat phase, **bottom row** – first entry into stimulus chamber post-rat exposure). **(B)** Mean firing rate of all assessment+, assessment-, flight+, and flight-cells identified in dPAG (**Top row** – pre-rat phase, **middle row** – post-rat phase, **bottom row** – first entry into stimulus chamber post-rat exposure).

## Supplementary Video Legends

***Movies S1-S3.*** Representative videos showing responses of assessment+ **(S1)**, assessment-**(S2)**, and flight+ **(S3)** cells recorded from dPAG.

***Movies S4-S6.*** Representative videos showing responses of assessment+ **(S4)**, flight+ **(S5)**, and flight-**(S6)** cells recorded from VMHdm.

## References

Amano K, Tanikawa T, Kawamura H, Iseki H, Notani M, Kawabatake H, Shiwaku T, Suda T, Demura H, Kitamura K (1982) Endorphins and pain relief. Further observations on electrical stimulation of the lateral part of the periaqueductal gray matter during rostral mesencephalic reticulotomy for pain relief. Appl Neurophysiol 45:123–135.

Arendt D, Tosches MA, Marlow H (2015) From nerve net to nerve ring, nerve cord and brain—evolution of the nervous system. Nat Rev Neurosci 17:61.

Bandler R, McCulloch T (1984) Afferents to a midbrain periaqueductal grey region involved in the “ defense reaction” in the cat as revealed by horseradish peroxidase. II. The diencephalon. Behav Brain Res 13:279–285.

Beitz AJ (1982) The organization of afferent projections to the midbrain periaqueductal gray of the rat. Neuroscience 7:133–159.

Beitz AJ (1989) Possible origin of glutamatergic projections to the midbrain periaqueductal gray and deep layer of the superior colliculus of the rat. Brain Res Bull 23:25–35.

Bergan JF, Ben-Shaul Y, Dulac C (2014) Sex-specific processing of social cues in the medial amygdala. eLife Sciences 3:e02743.

Blanchard DC, Li CI, Hubbard D, Markham CM, Yang M, Takahashi LK, Blanchard RJ (2003) Dorsal premammillary nucleus differentially modulates defensive behaviors induced by different threat stimuli in rats. Neurosci Lett 345:145–148.

Blanchard DC, Williams G, Lee EMC, Blanchard RJ (1981) Taming of wild Rattus norvegicus by lesions of the mesencephalic central gray. Psychobiology 9:157–163.

Braak H, Braak E (1992) Anatomy of the human hypothalamus (chiasmatic and tuberal region). Prog Brain Res 93:3–14; discussion 14–16.

Cameron AA, Khan IA, Westlund KN, Cliffer KD, Willis WD (1995) The efferent projections of the periaqueductal gray in the rat: A Phaseolus vulgaris-leucoagglutinin study. I. Ascending projections. J Comp Neurol 351:568–584.

Canteras NS (2002) The medial hypothalamic defensive system: hodological organization and functional implications. Pharmacol Biochem Behav 71:481–491.

Canteras NS, Chiavegatto S, Ribeiro do Valle LE, Swanson LW (1997) Severe reduction of rat defensive behavior to a predator by discrete hypothalamic chemical lesions. Brain Res Bull 44:297–305.

Canteras NS, Simerly RB, Swanson LW (1994) Organization of projections from the ventromedial nucleus of the hypothalamus: a Phaseolus vulgaris-leucoagglutinin study in the rat. J Comp Neurol 348:41–79.

Canteras NS, Simerly RB, Swanson LW (1995) Organization of projections from the medial nucleus of the amygdala: a PHAL study in the rat. J Comp Neurol 360:213–245.

Canteras NS, Swanson LW (1992) The dorsal premammillary nucleus: an unusual component of the mammillary body. Proc Natl Acad Sci U S A 89:10089–10093.

Choi GB, Dong H-W, Murphy AJ, Valenzuela DM, Yancopoulos GD, Swanson LW, Anderson DJ (2005) Lhx6 delineates a pathway mediating innate reproductive behaviors from the amygdala to the hypothalamus. Neuron 46:647–660.

Deng H, Xiao X, Wang Z (2016) Periaqueductal Gray Neuronal Activities Underlie Different Aspects of Defensive Behaviors. J Neurosci 36:7580–7588.

Falkner AL, Dollar P, Perona P, Anderson DJ, Lin D (2014) Decoding ventromedial hypothalamic neural activity during male mouse aggression. J Neurosci 34:5971–5984.

Falkner AL, Grosenick L, Davidson TJ, Deisseroth K, Lin D (2016) Hypothalamic control of male aggression-seeking behavior. Nat Neurosci 19:596–604.

Faturi CB, Rangel MJ Jr, Baldo MVC, Canteras NS (2014) Functional mapping of the circuits involved in the expression of contextual fear responses in socially defeated animals. Brain Struct Funct 219:931–946.

Fuchs SA, Edinger HM, Siegel A (1985) The organization of the hypothalamic pathways mediating affective defense behavior in the cat. Brain Res 330:77–92.

Graeff FG (1994) Neuroanatomy and neurotransmitter regulation of defensive behaviors and related emotions in mammals. Braz J Med Biol Res 27:811–829.

Gross CT, Canteras NS (2012) The many paths to fear. Nat Rev Neurosci 13:651–658.

Koutcherov Y, Mai JK, Ashwell KWS, Paxinos G (2002) Organization of human hypothalamus in fetal development. J Comp Neurol 446:301–324.

Kruk MR, van der Poel AM, de Vos-Frerichs TP (1979) The induction of aggressive behaviour by electrical stimulation in the hypothalamus of male rats. Behaviour 70:292–322.

Kunwar PS, Zelikowsky M, Remedios R, Cai H, Yilmaz M, Meister M, Anderson DJ (2015) Ventromedial hypothalamic neurons control a defensive emotion state. Elife 4 Available at: http://dx.doi.org/10.7554/eLife.06633.

Kurrasch DM, Cheung CC, Lee FY, Tran PV, Hata K, Ingraham HA (2007) The Neonatal Ventromedial Hypothalamus Transcriptome Reveals Novel Markers with Spatially Distinct Patterning. J Neurosci 27:13624–13634.

Lein ES et al. (2007) Genome-wide atlas of gene expression in the adult mouse brain. Nature 445:168–176.

Li C-I, Maglinao TL, Takahashi LK (2004) Medial amygdala modulation of predator odor-induced unconditioned fear in the rat. Behav Neurosci 118:324–332.

Lin D, Boyle MP, Dollar P, Lee H, Lein ES, Perona P, Anderson DJ (2011) Functional identification of an aggression locus in the mouse hypothalamus. Nature 470:221–226.

Malsbury CW, Kow L-M, Pfaff DW (1977) Effects of medial hypothalamic lesions on the lordosis response and other behaviors in female golden hamsters. Physiol Behav 19:223–237.

Mantyh PW (1982) Forebrain projections to the periaqueductal gray in the monkey, with observations in the cat and rat. J Comp Neurol 206:146–158.

Mantyh PW (1983) Connections of midbrain periaqueductal gray in the monkey. I. Ascending efferent projections. J Neurophysiol 49:567–581.

Martinez RC, Carvalho-Netto EF, Ribeiro-Barbosa ER, Baldo MVC, Canteras NS (2011) Amygdalar roles during exposure to a live predator and to a predator-associated context. Neuroscience 172:314–328.

Meller ST, Dennis BJ (1991) Efferent projections of the periaqueductal gray in the rabbit. Neuroscience 40:191–216.

Motta SC, Goto M, Gouveia FV, Baldo MVC, Canteras NS, Swanson LW (2009) Dissecting the brain ‘s fear system reveals the hypothalamus is critical for responding in subordinate conspecific intruders. Proc Natl Acad Sci U S A 106:4870–4875.

Nomoto K, Lima SQ (2015) Enhanced Male-Evoked Responses in the ventromedial hypothalamus of sexsually receptive female mice. Current Biology 25:589–594

Petrovich GD, Risold PY, Swanson LW (1996) Organization of projections from the basomedial nucleus of the amygdala: a PHAL study in the rat. J Comp Neurol 374:387–420.

Pfaff DW, Sakuma Y (1979a) Facilitation of the lordosis reflex of female rats from the ventromedial nucleus of the hypothalamus. J Physiol 288:189–202.

Pfaff DW, Sakuma Y (1979b) Deficit in the lordosis reflex of female rats caused by lesions in the ventromedial nucleus of the hypothalamus. J Physiol 288:203–210.

Remedios R, Kennedy A, Zelikowsky M, Grewe BF, Schnitzer MJ, Anderson DJ (2017) Social behaviour shapes hypothalamic neural ensemble representations of conspecific sex. Nature 550:388–392.

Schenberg LC, Póvoa RMF, Costa ALP, Caldellas AV, Tufik S, Bittencourt AS (2005) Functional specializations within the tectum defense systems of the rat. Neurosci Biobehav Rev 29:1279–1298.

Silva BA, Gross CT, Gräff J (2016a) The neural circuits of innate fear: detection, integration, action, and memorization. Learn Mem 23:544–555.

Silva BA, Mattucci C, Krzywkowski P, Cuozzo R, Carbonari L, Gross CT (2016b) The ventromedial hypothalamus mediates predator fear memory. Eur J Neurosci 43:1431–1439.

Silva BA, Mattucci C, Krzywkowski P, Murana E, Illarionova A, Grinevich V, Canteras NS, Ragozzino D, Gross CT (2013) Independent hypothalamic circuits for social and predator fear. Nat Neurosci 16:1731–1733.

Swanson LW (2000) Cerebral hemisphere regulation of motivated behavior. Brain Res 886:113–164.

Swanson LW, Petrovich GD (1998) What is the amygdala? Trends Neurosci 21:323–331.

Tessmar-Raible K, Raible F, Christodoulou F, Guy K, Rembold M, Hausen H, Arendt D (2007) Conserved Sensory-Neurosecretory Cell Types in Annelid and Fish Forebrain: Insights into Hypothalamus Evolution. Cell 129:1389–1400.

Tovote P, Esposito MS, Botta P, Chaudun F, Fadok JP, Markovic M, Wolff SBE, Ramakrishnan C, Fenno L, Deisseroth K, Herry C, Arber S, Lüthi A (2016) Midbrain circuits for defensive behaviour. Nature 534:206–212.

Vianna DML, Brandão ML (2003) Anatomical connections of the periaqueductal gray: specific neural substrates for different kinds of fear. Braz J Med Biol Res 36:557–566.

Wang L, Chen IZ, Lin D (2015) Collateral pathways from the ventromedial hypothalamus mediate defensive behaviors. Neuron 85:1344–1358.

Wilent WB, Oh MY, Buetefisch CM, Bailes JE, Cantella D, Angle C, Whiting DM (2009) Induction of panic attack by stimulation of the ventromedial hypothalamus. J Neurosurg 112:1295–1298.

Yang CF, Chiang MC, Gray DC, Prabhakaran M, Alvarado M, Juntti SA, Unger EK, Wells JA, Shah NM (2013) Sexually dimorphic neurons in the ventromedial hypothalamus govern mating in both sexes and aggression in males. Cell 153:896–909.

