## Supplementary figures and images for "Differential encoding of predator fear in the ventromedial hypothalamus and periaqueductal grey"

### Supplementary Materials

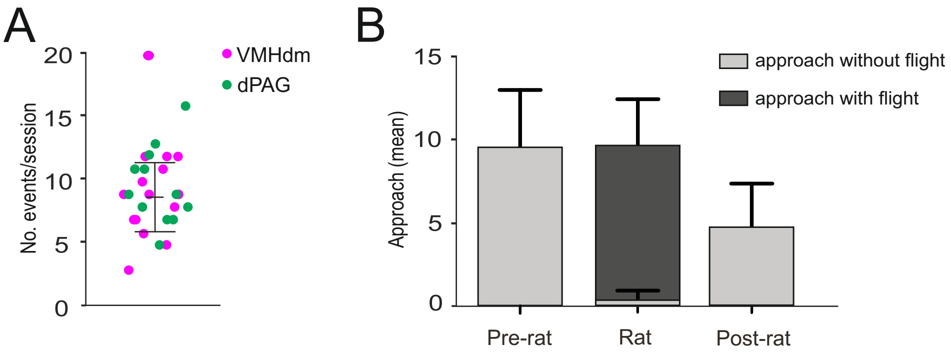

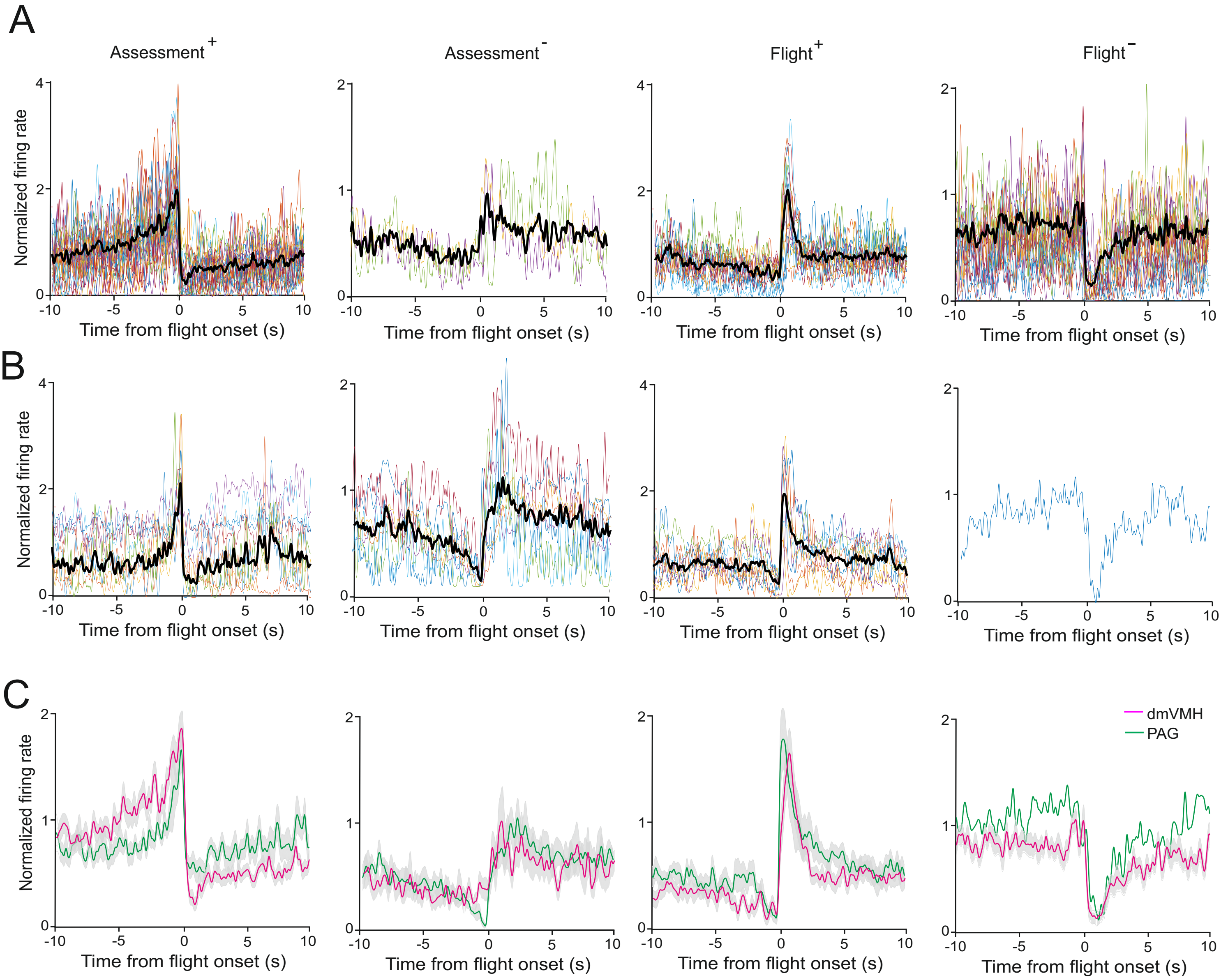

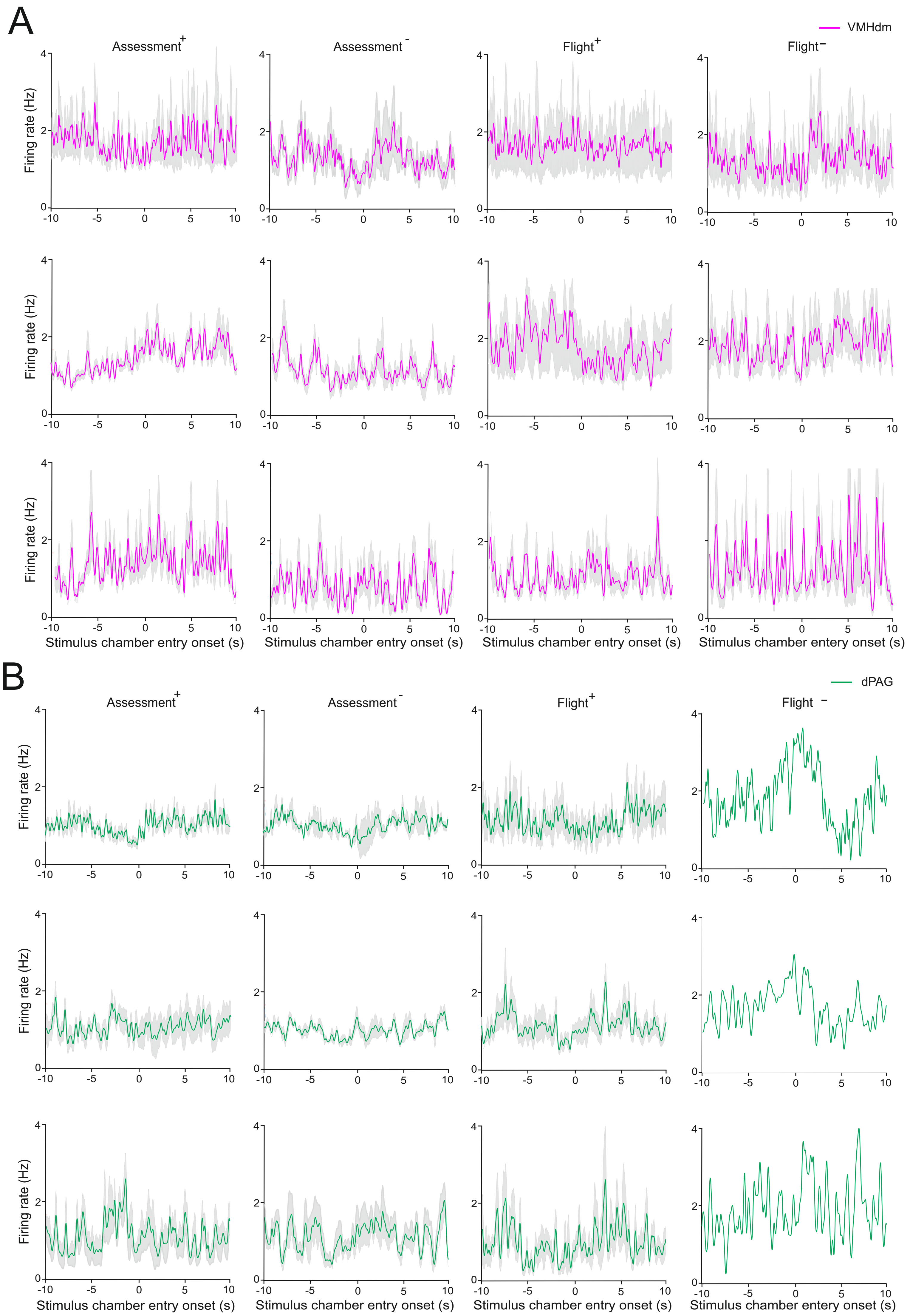
